# Discovery and genomic characterization of a novel wild clownfish hybrid from the Philippines

**DOI:** 10.64898/2026.07.14.738406

**Authors:** Lucy M. Fitzgerald, Floriane Coulmance, Théo Gaboriau, Anna Marcionetti, Sarah Schmid, Prince T. Apag, Annie G. Diola, Paul John Geraldino, Nicolas Salamin

## Abstract

Hybridization is widespread among marine fishes and can contribute to adaptation, diversification, and the generation of novel phenotypes. In clownfishes, only two wild hybrids *Amphiprion leucokranos* and *A. thiellei* have been described, yet those naturally occurring clownfish hybrids remain rarely documented. Both hybrids involve crosses with *A. sandaracinos*. During field surveys in the Philippines, we identified a previously undocumented clownfish individual with an unusual phenotype resembling both *A. sandaracinos* and *A. perideraion*. To characterize its origin, we combined genomic, mitochondrial, and phenotypic comparisons to previously described clownfish hybrid systems. Genome-wide PCA and admixture analyses supported mixed ancestry between *A. sandaracinos* and *A. perideraion*. Reconstruction of the mitochondrial genome placed the individual within the *A. sandaracinos* mitochondrial lineage. Together, these results support a hybrid origin and suggest predominant *A. sandaracinos* ancestry, consistent with a backcrossed descendant rather than a first-generation hybrid. Comparisons with the previously characterized hybrid *A. leucokranos* further revealed similarities in genomic composition and phenotype across independently derived clownfish hybrid systems. Our findings identify a previously undocumented natural clownfish hybrid and suggest that integrating genomic and field-based approaches may reveal additional cryptic hybrid systems and improve understanding of hybridization in clownfishes.

## Introduction

Hybridization is a common evolutionary process in marine systems that can contribute to adaptation, diver-sification, and the formation of novel phenotypes (Hobbs et al., 2022). Coral reef ecosystems contain many closely related species with overlapping geographic distributions, creating repeated opportunities for secondary contact and interspecific gene flow (Montanari et al., 2016). In the marine environment, hybridization is often concentrated in biogeographic contact regions or “suture zones” where historically isolated lineages come back into contact and exchange genetic material such as Christmas Island, Cocos (Keeling) Islands (Hobbs et al., 2009), and Socotra (DiBattista et al., 2015), where numerous hybrid reef fishes have been documented. The Philippines sits within the Indo-Malay-Philippine biodiversity hotspot, a region where Pacific and Indian Ocean lineages overlap and contribute to exceptional species richness, yet it is not recognized as a hybrid hotspot or suture zone (Gaither & Rocha, 2013). These systems provide insights into the ecological and evolutionary conditions that facilitate hybridization and shape species boundaries. Increasing evidence suggests that hybridization in marine fishes may be more prevalent than previously assumed, potentially reflecting the relatively weak reproductive barriers and broad dispersal capacities characteristic of many reef-associated species (Hobbs et al., 2022). However, hybrid systems may remain difficult to detect when hybrid individuals occur at low frequencies, occupy geographically restricted contact zones, or resemble parental species through introgression and backcrossing.

The frequency and evolutionary consequences of hybridization vary substantially among lineages depending on ecological interactions and reproductive behavior. Clownfishes, *Amphiprioninae*, provide an especially interesting system because their obligate association with host anemones strongly structures habitat use and species interactions. They are a monophyletic group of 28 species that are well known for their obligate mutualism with sea anemones that triggered their adaptive radiation (Litsios et al., 2012). Clownfish species differ in their degree of host specialization, ranging from generalists that use multiple anemone species to specialists restricted to one or two hosts. These ecological differences influence opportunities for interspecific encounters and hybridization. In addition, the overlapping appearances of many sympatric clownfishes can complicate identifying hybrid parental origins by phenotype alone. As a relatively recent radiation (10 mya) within damselfishes, clownfishes show evidence of hybridization both in natural populations and across their evolutionary history (Litsios & Salamin, 2014).

Despite their potential for hybridization, naturally occurring clownfish hybrids remain rare. To date, only two natural hybrids have been described: *Amphiprion leucokranos* and *A. thiellei. A. leucokranos*, a hybrid between *A. sandaracinos* and *A. chrysopterus*, has been extensively studied in Papua New Guinea and the Solomon Islands (Gainsford et al., 2015; Gainsford et al., 2020; Schmid et al., 2025). Originally described as a distinct species by Allen, 1973, subsequent ecological, morphological, and genetic analyses supported its status as a naturally occurring hybrid (Gainsford et al., 2015). More recent genomic analyses demonstrated extensive backcrossing, with almost all documented backcrosses occurring toward *A. sandaracinos* (Schmid et al., 2025), likely due to the size-based hierarchy (Gainsford et al., 2015). The second natural hybrid, *A. thiellei*, is thought to originate from hybridization between *A. sandaracinos* and *A. ocellaris*. However, this is purely based on circumstantial evidence of finding these species living together in cohabitation. *A. thiellei* was originally described in an aquarium magazine, based on a specimen reportedly collected near Cebu, Philippines (Burgess, 1981), yet no formal scientific study has examined its hybrid origin.

Both known clownfish hybrids share a common parental species of *A. sandaracinos* from the skunk complex (*akallopisos* group) which also includes *A. perideraion, A. akallopisos*, and *A. pacificus*. Previous studies have identified cytonuclear discordance and signatures of introgression within this group, suggesting that hybridization may have contributed to its evolutionary history (Litsios & Salamin, 2014; Marcionetti & Salamin, 2023). More recently, recurrent gene flow has been documented among members of the skunk complex (Marcionetti et al., 2024). In the Philippines, two species of the skunk complex *A. sandaracinos* and *A. perideraion* occur, but typically on different anemone species; *A. sandaracinos* is a *Stichodactyla* specialist and *A. perideraion* a *Radianthus* specialist (Gaboriau et al., 2025).

During fieldwork in the Philippines, we aimed to expand population-level sampling of clownfish species and investigate the possible occurrence of naturally occurring hybrids, particularly in the Cebu region where *A. thiellei* was originally reported. Based on phenotypic observations, we found a “hybrid-looking” individual cohabiting with *A. sandaracinos* individuals in the southern Philippines on the island of Mindanao and we tested the parentage using genomic analyses. Instead, we found a previously undocumented *A. sandaracinos* x *A. perideraion* hybrid. Interestingly, these two species differ in their host-specialization strategies making this hybridization event particularly surprising from an ecological perspective. This discovery also represents an example of a naturally occurring clownfish hybrid found outside a known hybrid hotspot, highlighting that hybridization in clownfishes may be more geographically widespread than previously recognized. We propose the name *Amphiprion x margaritokranos*, or “pearl-helmeted” clownfish, from the Greek “margarīťēs” (pearl) and “kranos” (helmeted).

## Methods

### Field Collection

On Talikud Island, Island Garden of Samal, Mindanao Philippines, we collected the hybrid sample (Figure 1A) during a SCUBA diving expedition where we were surveying local reefs. We collected genomic, ecological, and phenotypic data from *A. sandaracinos* and *A. perideraion* samples that were preserved in DESS buffer (Oosting et al., 2020). In total, we collected 11 new samples for this study: the hybrid individual, 6 *A. perideraion*, and 4 *A. sandaracinos* individuals from multiple sites in the Philippines (Table S1). These new *A. sandaracinos* and *A. perideraion* samples, together with previously published individuals from across their ranges, are shown in the distribution map in Figure 1B. Additional individuals from other clownfish species present on Talikud Island *A. biaculeatus, A. clarkii, A. frenatus, A. ocellaris*, and *A. polymnus*, were collected for preliminary parental species evaluation; these samples were used only in supplementary analyses (Table S2).

**Figure 1:**
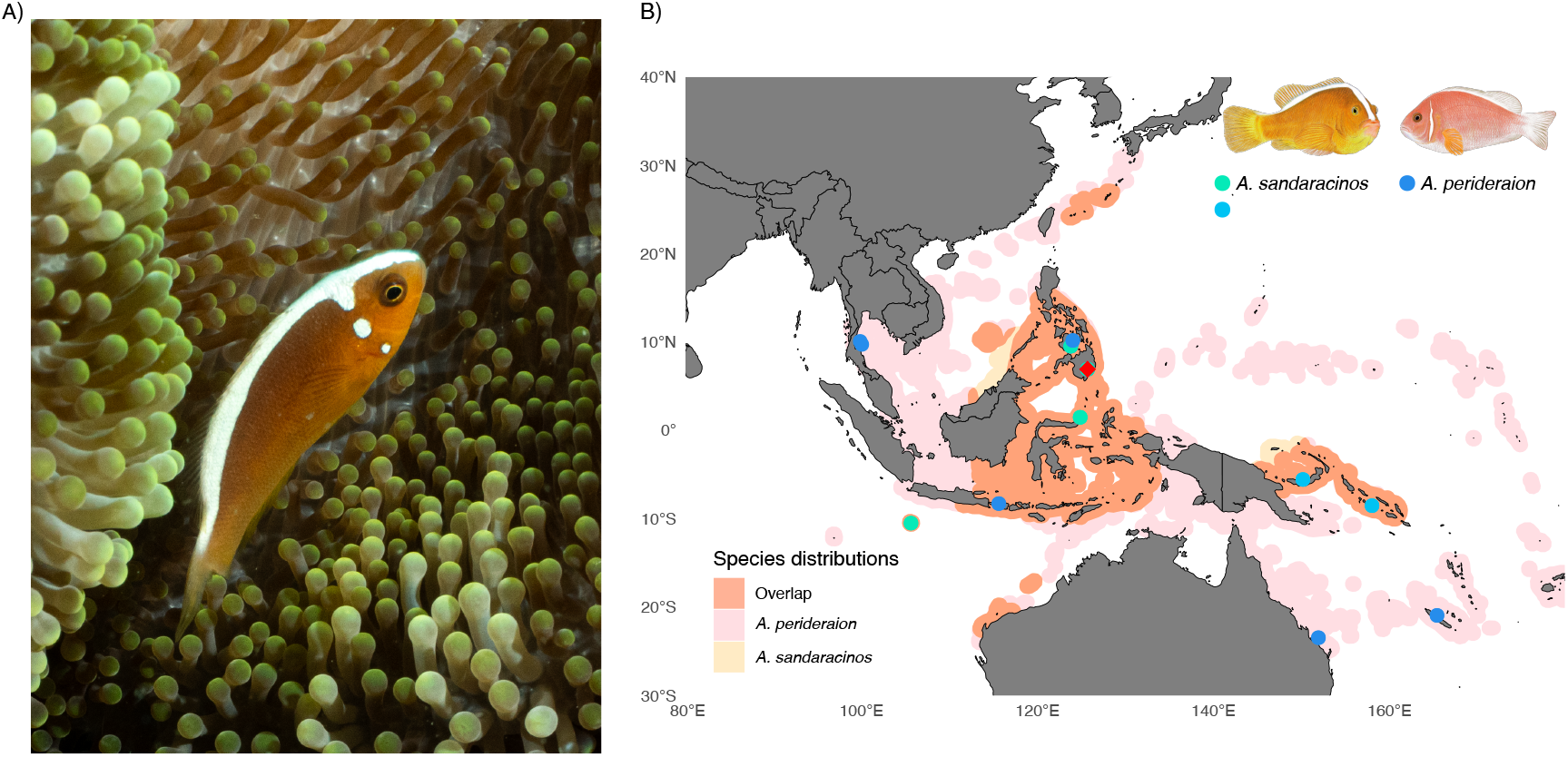
A) Photograph of the hybrid individual showing the intermediate phenotype between parental species *A. sandaracinos* x *A. perideraion* with two pearl-like dots on either side of its head, credit: Lucy Fitzgerald. B) Map of the species distribution and overlap for *A. sandaracinos* and *A. perideraion*. Sampling locations are shown as points: *A. sandaracinos* = green (central Indo-Pacific) and light blue (Bismark/Coral Sea), *A. perideraion* = dark blue, and the red diamond marks where the hybrid was sampled. The two *A. sandaracinos* colors represent the central Indo-Pacific and Bismarck/Coral Sea populations, which form two genetically divergent groups in our PCA and ADMIXTURE analyses.

### DNA extraction, library preparation and sequencing

DNA was extracted using DNeasy blood and Tissue Kit (Qiagen GmbH) and quantified using Qubit^®^ 2.0 Fluorometer (Thermo Fisher Scientific). DNA integrity was assessed by agarose gel electrophoresis. Libraries were prepared using the Nextera DNA Flex Library Preparation Kit following the manufacturer’s instructions, and fragment size distributions were validated using a Bioanalyzer (Agilent Technologies). Libraries were sequenced on the Element Biosciences Aviti platform using 150 bp paired-end reads at the Genomic Technologies Facility (GTF) of the University of Lausanne, Switzerland.

### Reads processing, mapping and SNP calling

Raw reads were trimmed and adapters removed using *Trimmomatic* v.0.39 (Bolger et al., 2014). Read quality was assessed using (*FastQC* v.0.11.9) and *MultiQC* (Ewels et al., 2016). Reads were mapped against the *A. ocellaris* reference genome (Ryu et al., 2022) using *BWA* v0.7.17 (Li & Durbin, 2009); and processed and sorted with *Samtools* v1.19.2 (Danecek et al., 2021). Mapping statistics were generated using *Bamtools* v2.5.2 (Barnett et al., 2011).

To identify the parental species of the hybrid, we first conducted preliminary analyses using only the newly collected samples from Talikud Island, including the hybrid. The additional clownfish species present at the site were *A. biaculeatus, A. clarkii, A. frenatus, A. ocellaris, A. perideraion, A. polymnus*, and *A. sandaracinos*. Based on the initial PCA and ADMIXTURE results, we identified *A. perideraion* as the most likely secondary parental species (Figure S1).

After confirming the parental species, we expanded the dataset by incorporating previously published *A. sandaracinos* and *A. perideraion* samples from across the Indo-Pacific (García-Jiménez et al., 2025; Marcionetti et al., 2024) to place the hybrid within a broader population genomic and geographic context. In addition, we included individuals from the other known hybrid system involving *A. sandaracinos, A. leucokranos* (*A. sandaracinos* x *A. chrysopterus*) using data from Schmid et al., 2025. These samples provided a comparative framework for interpreting ancestry proportions and hybrid classes in a well-characterized clownfish hybrid system.

Haplotypes were called with *GATK* v4.5.0.0 (Van der Auwera & O’Connor, 2020), then the gVCF files were merged with (*Picard Tools* v3.1.1) and followed by joint genotyping with *GATK*. We used *VCFtools* v0.1.16 and kept only biallelic SNPs with quality above 30 (`-minQ 30`), minimum depth 5 (`-minDP 5`), maximum depth 60 (`-maxDP 60`), max missingness 0.8 (`-max-missing 0.8`) and a minor allele frequency of 0.01.

### PCA & Admixture

Independent SNPs were selected by linkage disequilibrium (LD) pruning using *PLINK* v1.9 (Purcell et al., 2007) (`--indep-pairwise 50 5 0.2`). PCA and admixture proportions were estimated using *ADMIXTURE* v1.3.0 (Alexander et al., 2009) with *K*-values ranging from 2-6. The *K*-value with the lowest cross-validation error (`--cv`) was selected (*K* = 4).

### Local ancestry inference

Local ancestry was inferred using *ELAI* v1.01 (Guan, 2014), which estimates ancestry along the genome using a two-layer hidden Markov model. We used the filtered SNP dataset and specified *A. perideraion* and *A. sandaracinos* as the two parental reference populations. Analyses were performed separately for all 24 chromosomes using two ancestral populations (`-C 2`) and 10 upper-layer clusters (`-c 10`). We ran 30 expectation–maximization iterations (`-s 30`) and explored several values for the number of generations since admixture (`-mg = 1, 2, 3, 5, 10`). Posterior ancestry dosage estimates were subsequently averaged across the different `-mg` settings to obtain final local ancestry estimates for each SNP.

Following Schmid et al. (2025), dosage values were interpreted on a 0–2 scale, where values near 0 or 2 indicate homozygous ancestry from one parental species and values near 1 indicate heterozygous ancestry. Using the estimated dosage of *A. perideraion* ancestry, we classified SNPs as homozygous *A. sandaracinos* (dosage ≤ 0.2), heterozygous (0.8−1.2), homozygous *A. perideraion* (≥ 1.8), or intermediate, following thresholds adapted from Seixas et al. (2018) and Schmid et al. (2025).

### Mitochondrial genome reconstruction

The mitochondrial genome of the hybrid individual was reconstructed using *MITObim* v1.9.1 (Hahn et al., 2013). Iterative baiting and mapping were performed using the `--quick` option with a published *A. sandaracinos* mitochondrial genome (GB206) as a seed reference (Marcionetti et al., 2024). FASTQ headers were reformatted to ensure compatibility with the legacy MIRA/mirabait workflow. The reconstructed mitochondrial genome was aligned with published mitochondrial genomes from members of the skunk complex using *MAFFT*, and a maximum-likelihood phylogeny was inferred using *IQ-TREE* (Minh et al., 2020) with 1000 ultrafast bootstrap replicates. Trees were visualized in iTOL (Minh et al., 2020). The complete mitochondrial genome of *A. ocellaris* (Ryu et al., 2022) was included as an outgroup.

## Results

### Population structure & Admixture analyses

On the first principal component (PC1; 19.65% of explained variance), *A. sandaracinos* separated from *A. perideraion, A. chrysopterus* and the hybrids, including *A. leucokranos* and our new hybrid individual. The second principal component (PC2; 10.64%) revealed additional structure within *A. sandaracinos* with the hybrid clustering near a subset of *A. sandaracinos* individuals and *A. leucokranos* backcrosses (Figure 2A).

**Figure 2:**
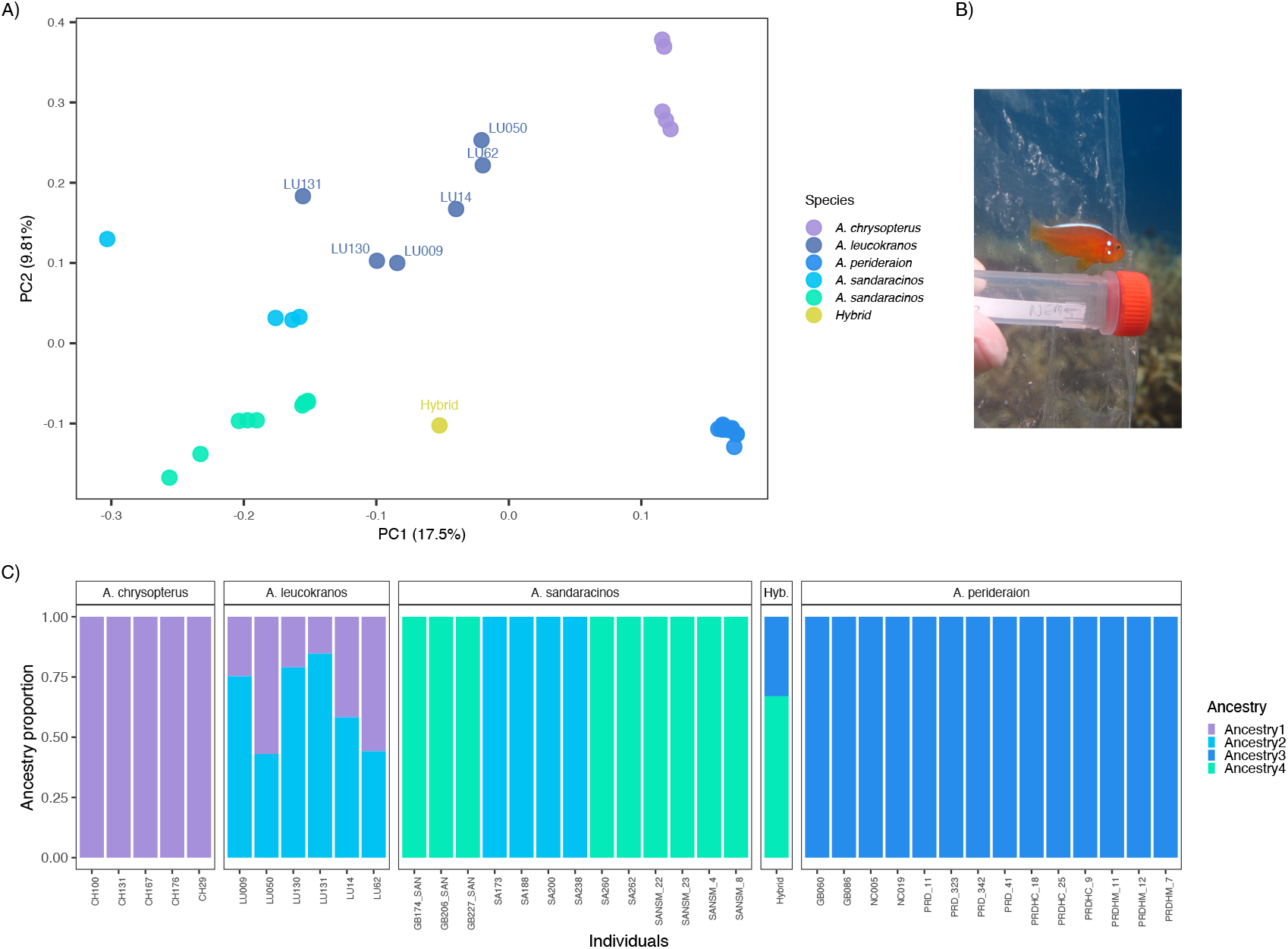
A) PCA with samples from *A. sandaracinos, A. perideraion, A. chrysopterus, A. leucokranos*, and the new hybrid, showing hybrids occupying intermediate positions between their parental species. The separation of *A. sandaracinos* into two clusters corresponds to the two geographically distinct populations shown in Figure 1. B) Photo of a first generation backcross *A. leucokranos* (LU130), credit: Ashton Gainsford. C) *ADMIXTURE* analysis with three ancestries corresponding to the three known clownfish species, with the two hybrids as admixed individuals of their parental species.

To estimate ancestry proportions, we used *ADMIXTURE*. The *K*-value with the lowest the crossvalidation, was *K* = 4 (CV=0.549) splitting *A. sandaracinos* into two ancestries, central Indo-Pacific and Bismark/Coral Sea (Figure 2). The *ADMIXTURE* analysis supports this interpretation with the hybrid exhibiting 30% *A. perideraion* ancestry which is consistent with a first-generation backcross rather than an equal contribution F1 hybrid. Its genomic profile closely parallels the ancestry proportions in previously characterized *A. leucokranos* backcrosses (e.g., LU009 and LU130) that exhibited 25% *A. chrysopterus* ancestry.

### Local ancestry along the genome

Local ancestry inference using *ELAI* revealed that the hybrid genome is dominated by heterozygous ancestry blocks between *A. sandaracinos* and *A. perideraion*, with a substantial portion of the remaining sites assigned to homozygous *A. sandaracinos* ancestry (Figure 3A). Homozygous *A. perideraion* segments were extremely rare, in line with what is expected for a backcross individual with the other parental species.

**Figure 3:**
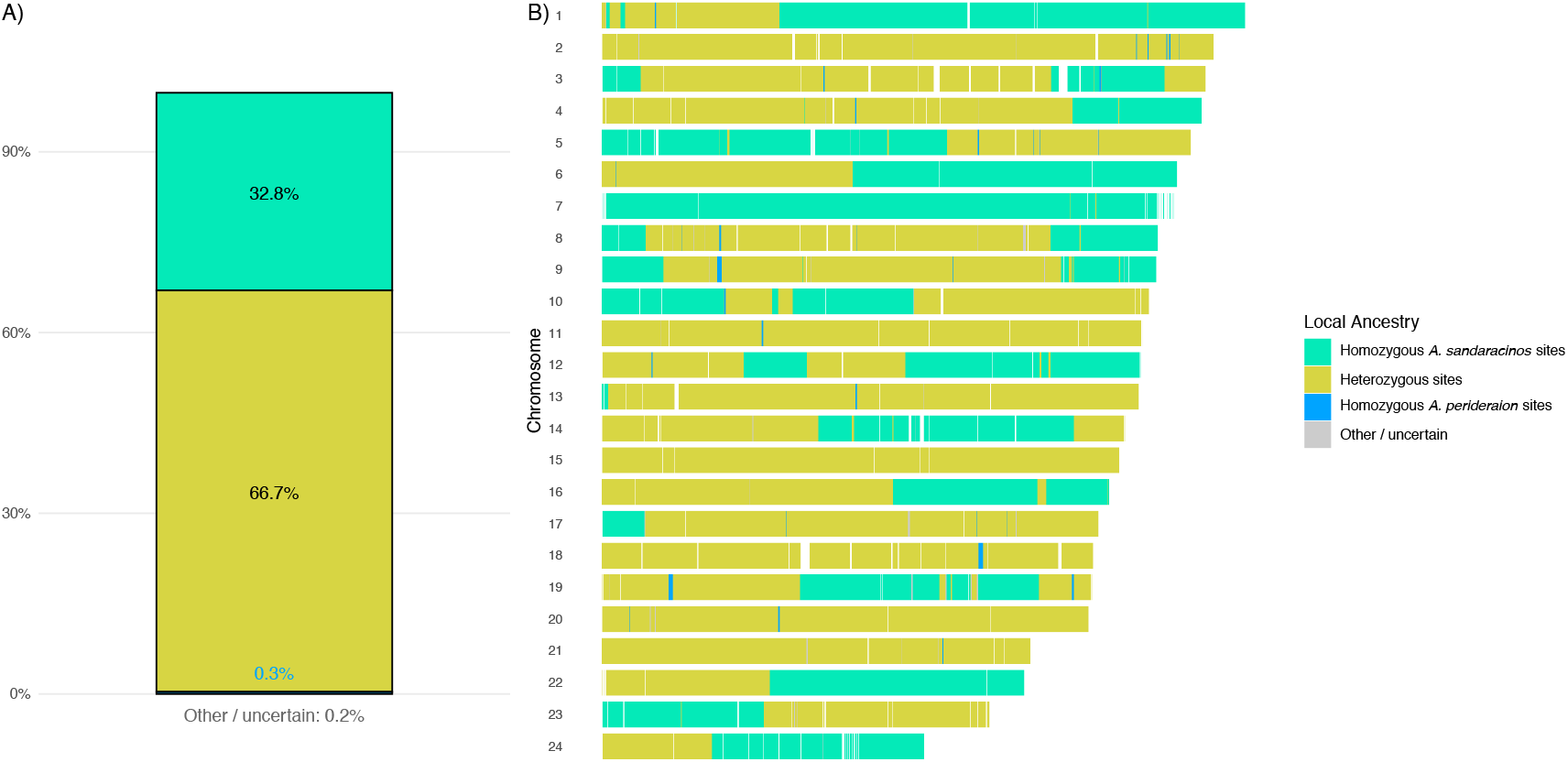
A) Genome-wide ancestry composition inferred using *ELAI* showing the relative proportions of homozygous *A. sandaracinos* (32.8%), heterozygous (66.7%), and homozygous *A. perideraion* (0.3%) ancestry states. B) Chromosome-level local ancestry plot displaying SNP-wise ancestry assignments across all 24 chromosomes. Each tile represents the inferred ancestry state at a single SNP, revealing a genome dominated by heterozygous ancestry blocks interspersed with *A. sandaracinos* homozygous regions and rare *A. perideraion* homozygous segments.

Chromosome-level patterns supported this ancestry composition. Most chromosomes contained long stretches of heterozygous ancestry interspersed with *A. sandaracinos* homozygous tracts, whereas *A. perideraion* homozygous blocks were short and infrequent (Figure 3B). This genomic mosaic closely resembles the structure observed in known *A. sandaracinos* backcrosses, including LU009 and LU130.

### Mitochondrial genome reconstruction

The reconstructed mitochondrial genome of the hybrid clustered within the *A. sandaracinos* mitochondrial lineage in the maximum-likelihood phylogeny (Figure S2). The hybrid grouped among *A. sandaracinos* individuals with strong bootstrap support and was clearly separated from *A. perideraion* and *A. akallopisos* mitochondrial lineages, indicating that it carries an *A. sandaracinos*-like mitochondrial genome.

Altogether, the PCA position, admixture proportions, ancestry structure, and mitochondrial placement consistently support a hybrid origin with predominant *A. sandaracinos* ancestry and maternal lineage. The resulting genomic mosaic closely mirrors the ancestry structure observed in previously characterized *A. leucokranos* backcrosses, reinforcing the conclusion that the individual represents a backcrossed descendant rather than an F1 hybrid.

## Discussion

We identified a novel naturally occurring clownfish hybrid that is phenotypically distinct from its parental species. This finding adds to the growing evidence that naturally occurring hybrid systems in clownfishes may be more diverse than currently recognized. Despite extensive overlap in species distributions and repeated opportunities for interspecific interactions, only two natural clownfish hybrids have been documented to date. The discovery of a previously undescribed hybrid in a region that has not been explored by researchers and aquarium trade raises the possibility that additional cryptic hybrid systems remain undetected but have been photographed by naturalists and divers.

Phenotypically, our hybrid resembles a combination of traits characteristic of both parental species including an intermediate body coloration and the presence of two small white “dots” on the head. This pattern is similar to the morphology of a previously documented *A. leucokranos* backcross (LU130) despite the distinct parental species in each hybrid system. The convergence in phenotype between these hybrid systems highlights how hybridization can generate similar intermediate traits even when genomic ancestries differ. At the same time, the comparison with *A. leucokranos* underscores that phenotype alone may not reliably reflect hybrid class, particularly because the individual we sampled was still juvenile and its adult pattern may differ slightly, reinforcing the importance of genomic data for resolving hybrid ancestry.

Our genomic analyses further support the hybrid origin of the individual. The PCA showed a clear separation among *A. sandaracinos, A. chrysopterus*, and *A. perideraion*, with hybrids occupying intermediate positions between parental species. Consistent with this pattern, the ADMIXTURE analysis indicated over half *A. sandaracinos* and less *A. perideraion* ancestry. This interpretation is strongly reinforced by the local ancestry results, which revealed a genome predominately composed of heterozygous ancestry blocks between *A. sandaracinos* and *A. perideraion*, with homozygous *A. sandaracinos* ancestry and rare homozygous *A. perideraion* segments. These proportions are what is expected for a first-generation backcross towards *A. sandaracinos*, rather than an equal-contribution F1 hybrid. When compared to the other hybrid system, the placement on the PCA and ancestry composition of our individual is broadly similar to early backcrosses *A. leucokranos* individuals (LU009 and LU130) which likewise exhibit predominately heterozygous genomes accompanied by a minority of homozygous *A. chrysopterus* blocks. These complementary lines of evidence demonstrate the value of integrating genome-wide clustering approaches when resolving hybrid class in natural systems.

The mitochondrial phylogeny further indicated that our hybrid carries an *A. sandaracinos* maternal lineage. Interestingly, Marcionetti et al., 2024 identified an Indonesian *A. sandaracinos* specimen (GB227) that clustered with *A. perideraion* in the mitochondrial phylogeny. This is in contrast to *A. leucokranos* where almost all the hybrids were found to be placed with *A. chrysopterus* on the mitochondrial tree despite being predominately backcrossed to *A. sandaracinos* (Schmid et al., 2025). Due to the limited sample size of only a single hybrid individual, additional individuals and population-level analyses will be necessary to determine the frequency of hybridization and resolve hybrid classes.

These contrasting mitochondrial patterns highlight how the size-based hierarchy in clownfish plays a role in the hybridization patterns and maternal lineages as stated by Gainsford et al., 2015. More generally, clownfishes present an unusual system in which ecological interactions, host specialization, and social structure may jointly influence opportunities for hybridization. Unlike many reef fishes, clownfishes occupy highly structured habitats defined by their host anemones and maintain strict size-based dominance hierarchies within social groups (Buston, 2003). These features influence both mate availability and the directionality of hybridization events. In addition, multiple clownfish species frequently cohabit the same anemone in the Coral Triangle, including the Philippines, increasing interspecific encounters and potentially facilitating hybridization (Camp et al., 2016). The occurrence of hybridization between species with different host specialization strategies further suggests that ecological boundaries between clownfish species may be more permeable than previously assumed. This is particularly interesting because host specialization has been proposed as an important axis of ecological divergence within clownfishes and has been associated with differences in morphology and ecology (Gaboriau et al., 2025).

We additionally observed geographic structuring within *A. sandaracinos* with individuals from the central Indo-Pacific clustering more closely with our hybrid, whereas samples from the Bismark/Coral Sea grouped more closely with *A. leucokranos*. This pattern is consistent with previous analyses by Schmid et al., 2025 and raises questions about the divergence of *A. sandaracinos* populations across its range. Such regional divergence within a single species can produce distinct ancestry signatures in genome-wide analyses, reflecting historical separation or limited gene flow between populations. This structuring may influence both the likelihood of hybridization and the genomic outcomes of hybridization events.

More broadly, our findings add to growing evidence that hybridization can shape the evolutionary history of coral reef fishes. Across marine systems, hybridization is increasingly recognized not simply as an evolutionary anomaly but as a recurrent process capable of influencing adaptation, generating phenotypic novelty, and contributing to patterns of diversification. Yet many natural hybrid systems remain difficult to identify because hybrids may occur at low frequencies, occupy geographically restricted regions, or become obscured through repeated backcrossing. In clownfishes specifically, accumulating evidence of contemporary hybridization together with signatures of historical introgression suggests that gene flow may have played a larger role in their evolutionary history than previously thought (Litsios & Salamin, 2014). Integrating field observations with genomic approaches may therefore reveal additional undocumented hybrid systems and improve understanding of the ecological and evolutionary processes generating diversity in coral reef communities.

## Supplementary Figures & Tables

**Table S1:** Sampling information for all individuals included in the main figures of this study. Hybrid status indicates previously characterized *A. leucokranos* hybrid classes from published datasets or the focal hybrid individual described in this study.

| Species | Sample ID | Country | Location | Citation | Hybrid status |
| --- | --- | --- | --- | --- | --- |
| *A. chrysopterus* | CH100 | Solomon Islands | Marovo Island | Schmid et al., 2025 | - |
| *A. chrysopterus* | CH131 | Solomon Islands | Marovo Island | Schmid et al., 2025 | - |
| *A. chrysopterus* | CH167 | Palau | Palau | Schmid et al., 2025 | - |
| *A. chrysopterus* | CH176 | Papua New Guinea | Kimbe Bay | Schmid et al., 2025 | - |
| *A. chrysopterus* | CH29 | Papua New Guinea | Kimbe Bay | Schmid et al., 2025 | - |
| *A. leucokranos* | LU009 | Papua New Guinea | Kimbe Bay | Schmid et al., 2025 | BC (1st gen) |
| *A. leucokranos* | LU050 | Papua New Guinea | Kimbe Bay | Schmid et al., 2025 | F1 |
| *A. leucokranos* | LU130 | Papua New Guinea | Kimbe Bay | Schmid et al., 2025 | BC (1st gen) |
| *A. leucokranos* | LU131 | Papua New Guinea | Kimbe Bay | Schmid et al., 2025 | BC (1st gen) |
| *A. leucokranos* | LU14 | Papua New Guinea | Kimbe Bay | Schmid et al., 2025 | F2 |
| *A. leucokranos* | LU62 | Papua New Guinea | Kimbe Bay | Schmid et al., 2025 | F1 |
| *A. perideraion* | GB060 | Indonesia | Tulamben | Marcionetti et al., 2024 | - |
| *A. perideraion* | GB086 | Indonesia | Tulamben | Marcionetti et al., 2024 | - |
| *A. perideraion* | NC005 | New Caledonia | Tibarama | Marcionetti et al., 2024 | - |
| *A. perideraion* | NC019 | New Caledonia | Tibarama | Marcionetti et al., 2024 | - |
| *A. perideraion* | PRD_11 | Thailand | Koh Tao | García-Jiménez et al., 2025 | - |
| *A. perideraion* | PRD_323 | Australia | Heron Island | García-Jiménez et al., 2025 | - |
| *A. perideraion* | PRD_342 | Australia | Heron Island | García-Jiménez et al., 2025 | - |
| *A. perideraion* | PRD_41 | Thailand | Koh Phangan | García-Jiménez et al., 2025 | - |
| *A. perideraion* | PRDHC_18 | Philippines | Talicut Island | This paper | - |
| *A. perideraion* | PRDHC_25 | Philippines | Talicut Island | This paper | - |
| *A. perideraion* | PRDHC_9 | Philippines | Cebu | This paper | - |
| *A. perideraion* | PRDHM_11 | Philippines | Bohol | This paper | - |
| *A. perideraion* | PRDHM_12 | Philippines | Bohol | This paper | - |
| *A. perideraion* | PRDHM_7 | Philippines | Cebu | This paper | - |
| *A. sandaracinos* | GB174_SAN | Indonesia | Manado | Marcionetti et al., 2024 | - |
| *A. sandaracinos* | GB206_SAN | Indonesia | Manado | Marcionetti et al., 2024 | - |
| *A. sandaracinos* | GB227_SAN | Indonesia | Manado | Marcionetti et al., 2024 | - |
| *A. sandaracinos* | SA173 | Papua New Guinea | Kimbe Bay | Schmid et al., 2025 | - |
| *A. sandaracinos* | SA188 | Solomon Islands | Marovo Island | Schmid et al., 2025 | - |
| *A. sandaracinos* | SA200 | Solomon Islands | Marovo Island | Schmid et al., 2025 | - |
| *A. sandaracinos* | SA238 | Papua New Guinea | Kimbe Bay | Schmid et al., 2025 | - |
| *A. sandaracinos* | SA260 | Australia | Christmas Island | Schmid et al., 2025 | - |
| *A. sandaracinos* | SA262 | Australia | Christmas Island | Schmid et al., 2025 | - |
| *A. sandaracinos* | SANSM_22 | Philippines | Talicut Island | This paper | - |
| *A. sandaracinos* | SANSM_23 | Philippines | Talicut Island | This paper | - |
| *A. sandaracinos* | SANSM_4 | Philippines | Bohol | This paper | - |
| *A. sandaracinos* | SANSM_8 | Philippines | Bohol | This paper | - |
| Hybrid | Hybrid | Philippines | Talicut Island | This paper | BC (1st gen) |

**Figure S1:**
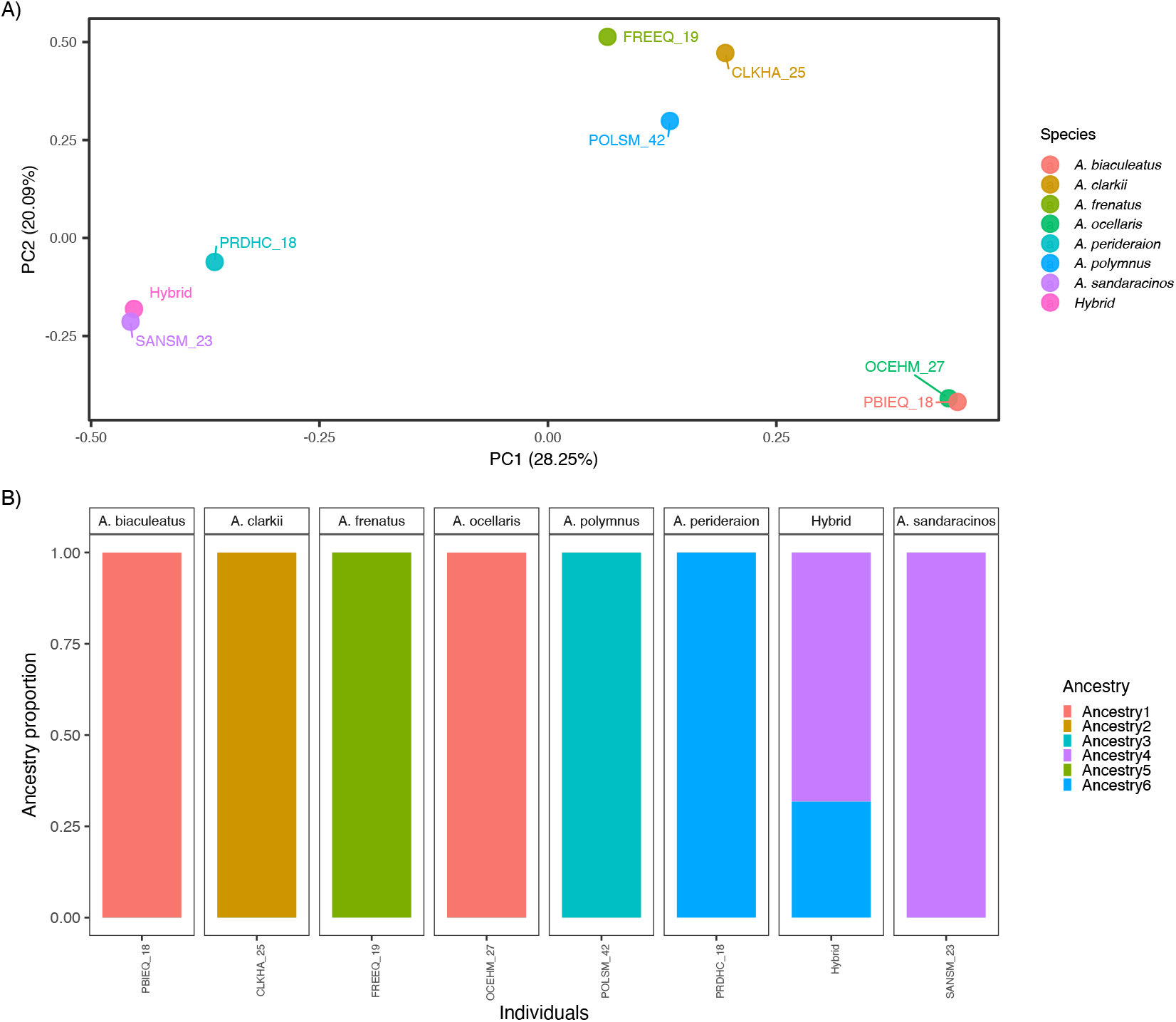
Population structure of potential parental species occurring on Talikud Island, Davao City, Mindanao, Philippines, and the hybrid individual. A) Principal component analysis (PCA) of genome-wide SNP variation among the sampled individuals. Each point represents an individual, colors indicate species identity, and labels correspond to individual IDs. B) ADMIXTURE analysis showing individual ancestry proportions inferred from genome-wide SNP data. Together, these analyses were used to assess the genetic relationships among species present on Talikud Island and to identify the most likely parental taxa contributing to the hybrid individual.

**Figure S2:**
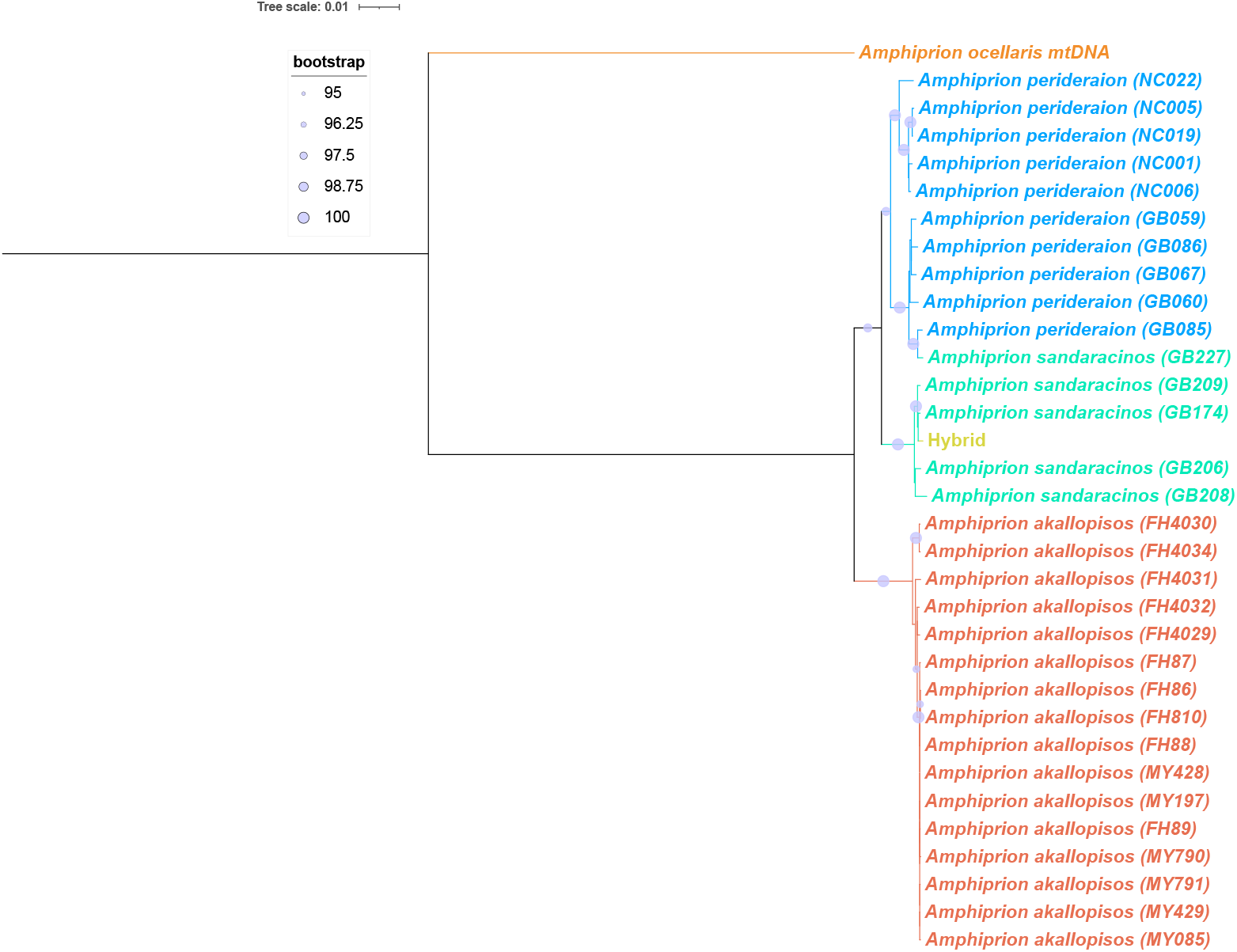
Maximum-likelihood mitochondrial phylogeny inferred from complete mitochondrial genome sequences of members of the skunk complex (*A. sandaracinos, A. perideraion*, and *A. akallopisos*) including the hybrid individual. Bootstrap support values are shown at internal nodes. The reconstructed hybrid mitochondrial genome clusters within the *A. sandaracinos* lineage.

**Table S2:** Sampling information for additional individuals included in supplementary analyses. These samples were used either for the full species comparisons (Figure S1) or mitochondrial phylogeny (Figure S2)

| Species | Sample ID | Dataset type | Citation |
| --- | --- | --- | --- |
| *A. perideraion* | NC022 | Mito tree | Marcionetti et al., 2024 |
| *A. perideraion* | NC001 | Mito tree | Marcionetti et al., 2024 |
| *A. perideraion* | NC006 | Mito tree | Marcionetti et al., 2024 |
| *A. perideraion* | GB059 | Mito tree | Marcionetti et al., 2024 |
| *A. perideraion* | GB067 | Mito tree | Marcionetti et al., 2024 |
| *A. perideraion* | GB085 | Mito tree | Marcionetti et al., 2024 |
| *A. sandaracinos* | GB208 | Mito tree | Marcionetti et al., 2024 |
| *A. sandaracinos* | GB209 | Mito tree | Marcionetti et al., 2024 |
| *A. akallopisos* | FH4030 | Mito tree | Marcionetti et al., 2024 |
| *A. akallopisos* | FH4034 | Mito tree | Marcionetti et al., 2024 |
| *A. akallopisos* | FH4031 | Mito tree | Marcionetti et al., 2024 |
| *A. akallopisos* | FH4032 | Mito tree | Marcionetti et al., 2024 |
| *A. akallopisos* | FH4029 | Mito tree | Marcionetti et al., 2024 |
| *A. akallopisos* | FH86 | Mito tree | Marcionetti et al., 2024 |
| *A. akallopisos* | FH810 | Mito tree | Marcionetti et al., 2024 |
| *A. akallopisos* | FH87 | Mito tree | Marcionetti et al., 2024 |
| *A. akallopisos* | FH88 | Mito tree | Marcionetti et al., 2024 |
| *A. akallopisos* | MY428 | Mito tree | Marcionetti et al., 2024 |
| *A. akallopisos* | MY197 | Mito tree | Marcionetti et al., 2024 |
| *A. akallopisos* | FH89 | Mito tree | Marcionetti et al., 2024 |
| *A. akallopisos* | MY791 | Mito tree | Marcionetti et al., 2024 |
| *A. akallopisos* | MY085 | Mito tree | Marcionetti et al., 2024 |
| *A. akallopisos* | MY790 | Mito tree | Marcionetti et al., 2024 |
| *A. akallopisos* | MY429 | Mito tree | Marcionetti et al., 2024 |
| *A. biaculeatus* | PBIEQ_18 | All species analysis | This paper |
| *A. clarkii* | CLKHA_25 | All species analysis | This paper |
| *A. frenatus* | FREEQ_19 | All species analysis | This paper |
| *A. ocellaris* | OCEHM_27 | All species analysis | This paper |
| *A. polymnus* | POLSM_42 | All species analysis | This paper |

## Declarations

LMF and NS conceptualized and designed the study. LMF conducted the research, performed the analyses, interpreted the results, and wrote the first draft of the manuscript. Fieldwork expeditions and sample collection were carried out by LMF, FC, TG, and PTA. AM and SS contributed to the genomic analyses and interpretation of the results. All authors reviewed, revised, and approved the final version of the manuscript.

## Acknowledgments

We would like to thank Laurie Mitchell for their comments on this manuscript. Additionally, the South Shore Divers especially Angel Santa Cruz for their help in coordinating the diving in Davao City, Mindanao, Philippines. We thank the DCSR infrastructure of the University of Lausanne for the computing resources and the USC Department of Biology for their help in obtaining permits. Lastly, Athina Gavriilidou in the Department of Computational Biology for the Greek name.

## Data Accessibility Statement

Data and scripts will be available on Github. Genomic data in the form of raw reads will be deposited in the NCBI Sequence Read Archive (SRA) under the bioproject ID:XX. Access to these data will also be provided upon acceptance of the manuscript.

### Benefits-Sharing Statement

The sharing of our data and results on public databases ensures transparency and facilitates further research in the field. All fieldwork conducted during this study adhered to local regulations and was carried out in collaboration with local entities. Our research activities were conducted in accordance with the “Access and Benefit Sharing” (ABS) principles of the Nagoya Protocol established by the Convention on Biological Diversity (CBD).

Samples from the Philippines were obtained by Nicolas Salamin’s research group in collaboration with the University of San Carlos (PTA, AD, and PJG contributed to obtaining permits, sampling, and DNA extraction).

### Biological Sampling Permits

The research permits obtained for this study: Philippines Gratuitous Permit from BFAR Region XI. Permit No 03025-032625.

